# A Method for Preparing Morphologically Preserved Wildlife Fecal Specimens for Long-Term Ecological Studies

**DOI:** 10.1101/2025.04.15.649043

**Authors:** Zhang Jiahao, Zhang Dongling, Xu Xinrui, Zhang Yunqiao, Dai Qiang

## Abstract

1. Wildlife feces are a valuable noninvasive resource in ecological and conservation research. However, traditional preservation methods are unable to maintain morphological integrity while simultaneously preserving the biological and chemical composition of fecal samples.
2. This study presents a novel method for the preparation of fecal specimens through a multistep immersion process using sodium carboxymethyl cellulose, sodium benzoate, clotrimazole, ethanol, pyrethroid emulsion and polyvinylpyrrolidone solution.
3. The specimens produced by this method exhibited high mechanical strength, ensuring durability and resistance to handling damage. During a storage period of six months, this method successfully preserved the morphological characteristics of fecal samples while maintaining DNA integrity, with no signs of mold or insect damage. DNA extraction achieved a 100% success rate and species identification remained consistent with fresh samples, with BLAST match rates exceeding 99% in 15 specimens. In addition, heavy metals such as chromium, arsenic, and lead were detected in fecal samples from different species.
4. By allowing long-term preservation of fecal samples, this method transforms urine from a short-term diagnostic tool into a durable resource for monitoring biodiversity. It can extend the applications of fecal samples across spatial and temporal scales, reinforcing their role in ecological research and global conservation of biodiversity.

## Introduction

As noninvasive biological samples, wildlife feces have played a significant role in ecological and conservation biology research (Mondol et al. 2009; Węgrzyn et al. 2018; Navarro-Castilla et al. 2023). Since Seton (1925) introduced ‘scatology’ to mammalogy, fecal research has expanded across various scientific disciplines (Hartman et al. 1958). Fecal samples are particularly valuable in field surveys where direct observation or population counts are difficult, especially for elusive or rare species (Palomares et al. 2002). Modern scatological studies address a wide range of fundamental ecological questions, including animal behavior biodiversity (Boyer et al. 2015), habitat (Bearer et al. 2008), and populations (Monterroso et al. 2019). Additionally, urine also provides valuable information for disciplines like behavior (Navarro-Castilla et al. 2019), genetics (Oja et al. 2017), and physiology (Barja et al. 2007).

Despite the value of fecal analysis, accurate fecal identification remains a significant challenge, particularly in regions with high species diversity (Hawlitschek et al. 2018). Misidentifications of feces can lead to alarmingly erroneous outcomes, raising concerns about the reliability of feces-based field surveys (Spitzer et al. 2019). Many species produce morphologically similar feces, and a single species, or even a single individual, fecal characteristics can vary due to dietary differences (Magondu et al. 2023), making direct identification in the field more challenging. Although specialized training enhances accuracy, ensuring that all survey participants have sufficient skills for accurate fecal identification remains a challenge. This problem extends beyond volunteers and rangers to professional researchers, who are also prone to errors (Costa et al. 2017).

Collecting and preparing biological specimens are essential for science and conservation (Rocha et al. 2014; He et al. 2021; Nachman et al. 2023). Feces serve as an important noninvasive resource in wildlife biology and ecological research (Biswas et al. 2019), yet effective methods for their long-term preservation remain lacking. Current fecal sampling and storage protocols are primarily aimed at DNA extraction and analysis, including frozen storage (Shores et al. 2015), dry sampling with silica (Murphy et al. 2000), ethanol preservation(Murphy et al. 2002), and a two-step approach involving ethanol storage followed by silica desiccation(Roeder et al. 2004). Although these methods effectively preserve DNA (Biswas et al. 2019), they do not retain fecal morphology over extended periods. The inability to preserve biological and chemical composition and morphological information limits the broader application of fecal samples in ecological studies, particularly in retrospective analyses and comparative research.

We believe that fecal specimens that preserve both morphology and biological and chemical composition offer significant potential for ecological and conservation research. These specimens, especially those with species confirmation by DNA analysis, can serve as valuable training materials for investigators, providing a more direct and reliable reference than photographs or descriptions, thus reducing misclassification errors. Furthermore, preserved fecal specimens act as verifiable records of field surveys, supporting data validation, and enhancing the repeatability and reliability of ecological studies. Beyond their immediate research applications, fecal specimens also hold promise as long-term records of ecological processes, offering insights into environmental degradation (Malaney & Cook 2018), pollutant exposure (Tran et al. 2015), dietary shifts(Blumenthal et al. 2012), pathogen transmission (Delahoy et al. 2018), and other areas of study as research methods evolve. This is similar to how museum specimens are now used to study historical ecological changes (Wandeler et al. 2007; Staats et al. 2013; Burrell et al. 2015). Additionally, thoroughly sterilized fecal specimens can serve as valuable resources in nature education, providing an interactive way to enhance learning through direct, hands-on experiences.

This study presents an innovative approach for preparing mammalian fecal specimens that preserves morphology and biological and chemical composition, providing a reliable foundation for verification and future analyzes. To evaluate the effectiveness of this method, we applied it to fecal samples of various carnivores and herbivores, evaluating their morphology, heavy metal content, and DNA Quality after six months of preservation. This approach not only ensures the long-term preservation of fecal specimens, but also enhances their potential for ecological and conservation research, particularly in the context of global environmental change.

## Materials and Methods

### Field Collecting

Fecal samples were collected from the forest in the Yingjing area of Giant Panda National Park, China. Intact feces were selected during field collection, and their original morphology was photographed for documentation prior to placement in sampling boxes. A layer of drying silica gel was added to the bottom of the container, followed by the fecal sample arranged to maintain its original shape and completely covered with additional silica gel. To minimize damage during transport, a layer of acrylic fiber or sterile cotton was placed on top of silica gel for cushioning. The sampling boxes were sealed with lids and then wrapped in cling film to prevent dispersal of contents and protect against the ingress of external moisture and contaminants. Fresh samples should be prioritized if fecal samples are intended for DNA extraction. The samples were transported with careful attention to dryness, and the drying silica gel was promptly replaced when saturated.

### Preparing Procedure

First, saturate the fecal sample with 4% sodium carboxymethylcellulose solution, followed by air drying in a cool place for 8 hours to fix the specimens. Depending on the size of the fecal sample, the saturation and drying steps may need to be repeated three or more times to ensure thorough fixation. The fecal specimens were then saturated with a series of ethanol solutions (75%, 85%, 95% and 100%), followed by air drying for 2 to 4 h after each step. Then 0.1% benzoate ethanol solutions and 0.5% clotrimazole acetone solutions were used for antibacterial and antifungal treatment. The fecal specimens were saturated with each solution at least three times, with air drying for 8 hours following each saturation. An additional overnight air drying after those treatments was performed to ensure complete evaporation of the acetone. Subsequently, the specimens were soaked in a 30% polyvinylpyrrolidone (PVP) solution to reinforce their structure, followed by air drying for 1 to 2 days. The fecal specimens were treated with anti-insects with a 2.5% pyrethroid emulsion for 3 minutes, followed by drying in a fume hood for 4 hours. Finally, the fecal specimens were placed in a drying oven at 30 ° C (Figure 1).

**FIGURE 1.**
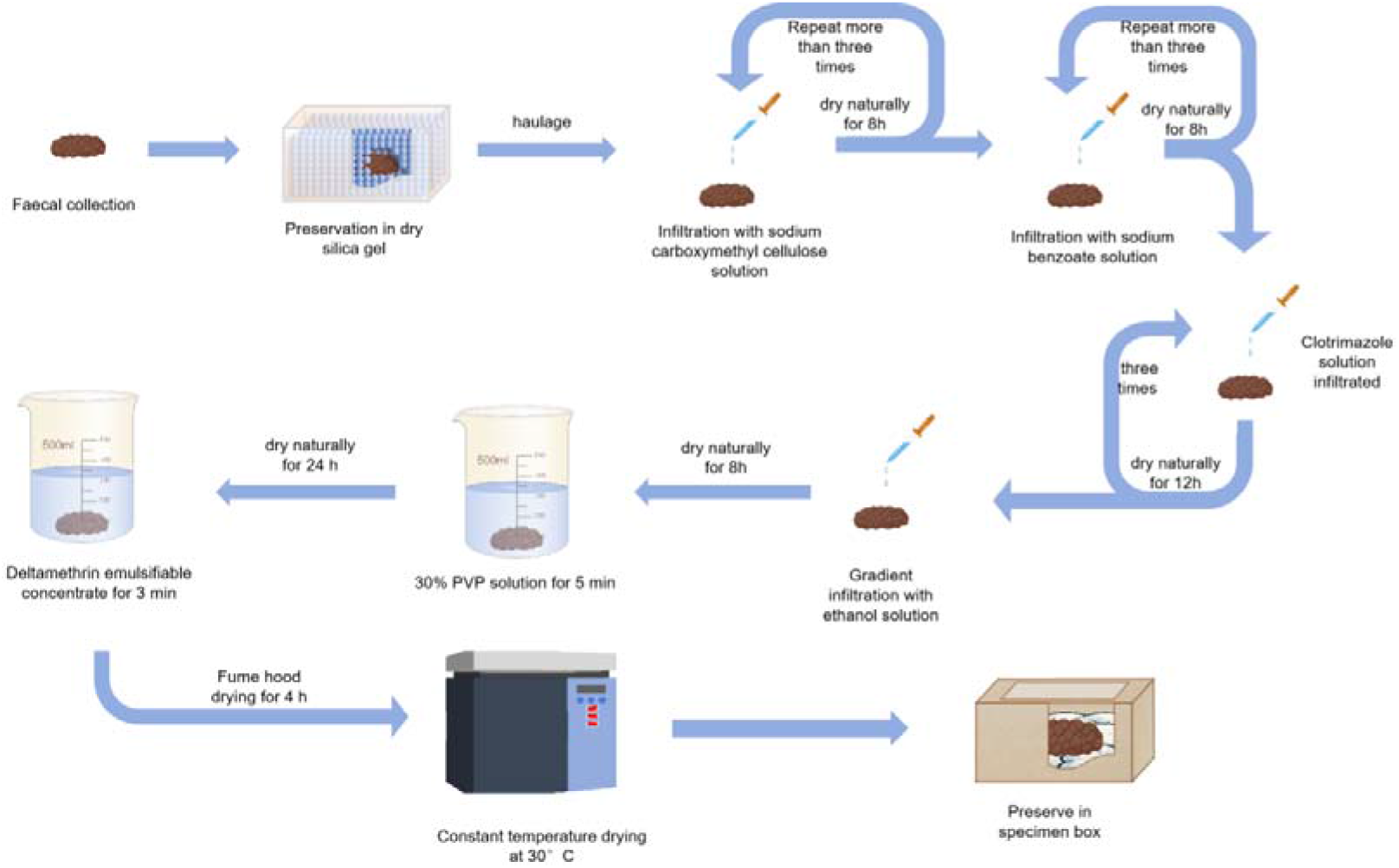
Workflow for Fecal Sample Reparation.

### Specimen Preservation

To prevent insect damage, one to two mothballs were placed at the bottom of the sample box, followed by a layer of acrylic fiber and some color-changing silica gel. The fecal specimens were then placed on top (Figure 1). Finally, the specimens were labeled, recorded, and archived according to museum protocols.

### Mechanical Properties

Shore hardness was measured using an LX-A dual-needle Shore durometer (GOYOJO Industrial Technology, Shenzhen, China) under static compression conditions, with five measurements taken per specimen. This test evaluates a material’s resistance to indentation and is widely used to assess the mechanical properties of polymers, elastomers, and biological specimens (Falanga & Bucalo 1993; Basfar 1997). The LX-A durometer, designed for soft materials, applies a standardized force to determine the firmness and structural integrity of fecal specimens. Hardness values range from 0 (extremely soft, gel-like materials) to 100 (hard rubber-like materials), providing insights into the durability of preserved specimens and ensuring their suitability for long-term storage and handling.

### DNA Quality Assessment

A 220-260 mg sample was carefully scraped from the surface of each feces using a sterile scalpel and transferred to a 2 ml centrifuge tube for further processing. To avoid cross-contamination, forceps, scalpel, Petri dishes, and bench surfaces were immediately sanitized with ethanol or replaced after handling each sample. DNA was extracted with the QIAamp® Fast DNA Stool Mini Kit (Qiagen) according to the manufacturer’s instructions. The total DNA extracted was stored at -20 ° C until further use.

We used PCR amplification on two selected gene loci (Table 1). Each PCR reaction mixture contained 2 μl DNA template, 10 μl Taq PCR Master Mix (Shenggong Biological Engineering Co., Ltd.), 0.8 μl each forward and reverse primer, and 6.4 μl nuclease-free water (ddH2O), for a total volume of 20 μl. Specific PCR protocols were used for each gene locus. For 16S rDNA, the conditions were: initial denaturation at 95 ° C for 2.5 min, followed by 35 cycles of 95 ° C for 30 s, 50 ° C for 30 s, and 72 ° C for 1 min, with a final extension at 72 ° C for 10 min. For Cytb, the conditions were: initial denaturation at 95 ° C for 2.5 min, followed by 35 cycles of 95 ° C for 30 s, 55 ° C for 30 s, and 72 ° C for 1 min, with a final extension at 72 ° C for 10 min. All PCR products were stored at 4 ° C.

**Table 1.**
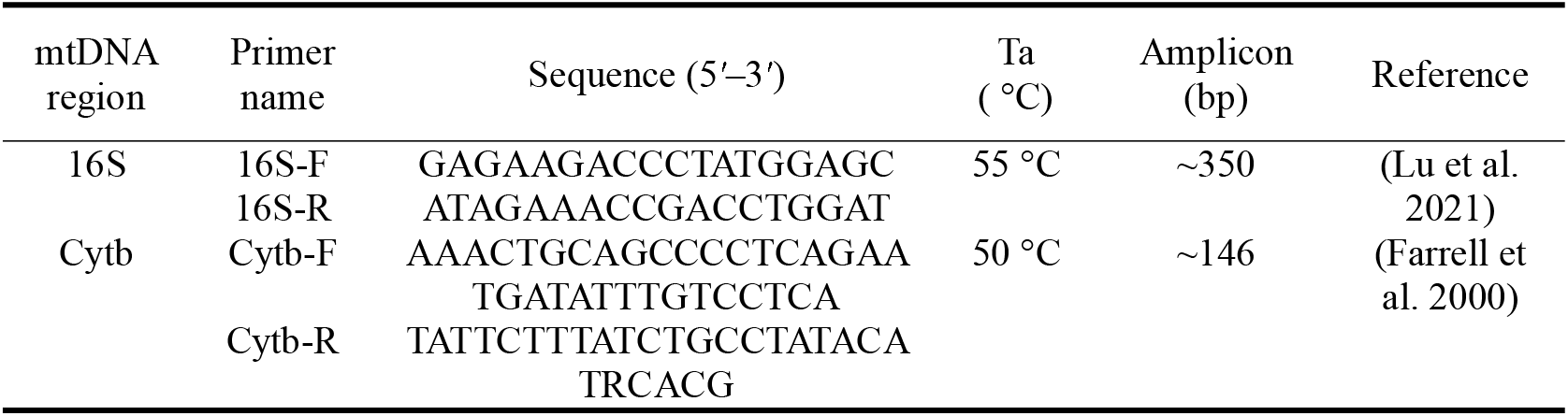
Primer sequences for mitochondrial genes.

**Table 2.** Shore hardness measurements of fecal specimens.

| Fecal Specimens | Hardness (Shore A) |  |  |
| --- | --- | --- | --- |
|  | Mean | Range | Standard Deviation |
| *Elaphodus cephalophus* | 84.92 | 81.4~89.9 | 3.03 |
| *Ailurus styani* | 36.02 | 25.6~47.1 | 9.08 |
| *Macaca thibetana* | 79.36 | 68.8~84.5 | 5.63 |
| *Hystrix brachyura* | 82.46 | 72.8~90.0 | 6.51 |
| *Sus scrofa* | 85.04 | 75.5~93.0 | 6.13 |
| *Prionailurus bengalensis* | 81.38 | 74.3~89.4 | 5.67 |

DNA was first extracted from fecal samples for species identification prior to the sample preservation procedure. After six months of storage, DNA was reextracted from the same samples to assess DNA quality after preservation. The assembled sequence was confirmed by a homology search in the GenBank database using BLAST. The identification of species was based on a 97% to 100% similarity match, with the best match selected as the species corresponding to the sample.

### Heavy-Metal Detection

The fecal samples were dried at 50 ° C, ground and sieved through a 0.15 mm metal sieve. 300 mg of fecal samples were digested in a 3:1 solution of HNO3 - HClO4 (v / v) to determine the concentrations of Cr, Cu, Zn, As, Cd, Pb, and Hg (Jin et al. 2023). After digestion, the samples were filtered and diluted to 50 ml with ultrapure water. Elemental concentrations were measured using inductively coupled plasma mass spectrometry (ICP-MS).

## Results

### Morphology and Hardness

Overall, the processed specimens exhibited high structural integrity and resistance to damage, effectively maintaining the original morphology of the feces. As shown in Figure 2, the specimens retained the color and sheen of fresh feces, with clearly visible animal hairs or plant fibers, indicating that key external characteristics were well preserved. Furthermore, during the storage period, no signs of decay, mold growth, or insect infestation were observed, demonstrating the effectiveness of the preservation method.

**FIGURE 2.**
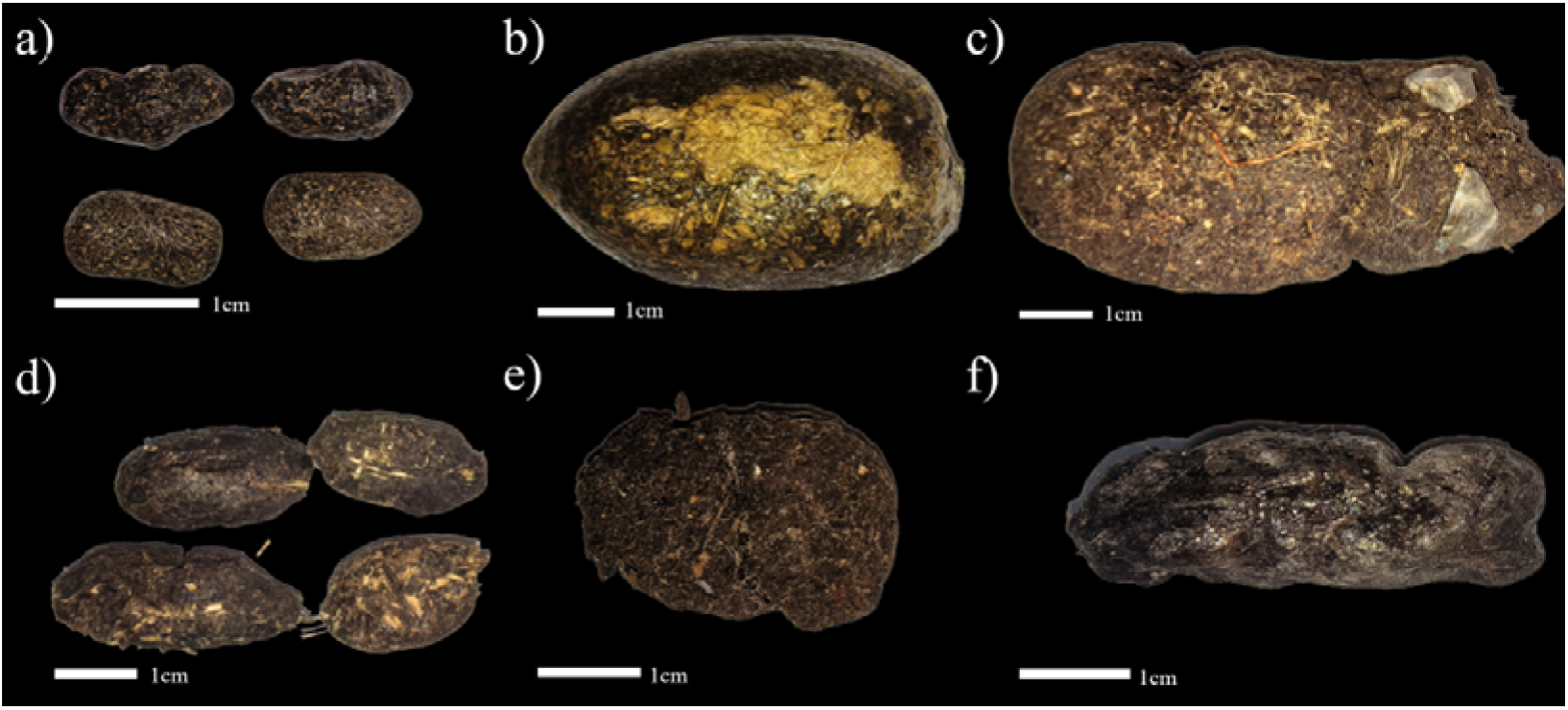
Photos of fecal specimens of (a) *Elaphodus cephalophus*, (b) *Ailurus styani*, (c) *Macaca thibetana*, (d) *Hystrix brachyura*, (e) *Sus scrofa* and (f) *Prionailurus bengalensis*.

Shore hardness tests revealed that most specimens exhibited hardness values between 79 and 85, comparable to firm rubber or dense leather, indicating high structural integrity. However, *Ailurus styani samples* had a significantly lower average hardness of 36.02, similar to soft rubber or an eraser, suggesting a much softer and less rigid structure.

### DNA quality

A total of six fecal samples were analyzed to assess DNA quality, with DNA extracted from fresh samples and after six months of preservation as samples. Gel electrophoresis was performed on all samples. As shown in Figure 3, DNA was successfully extracted from all preserved fecal samples, except Sample 6, which may have undergone degradation.

**FIGURE 3.**
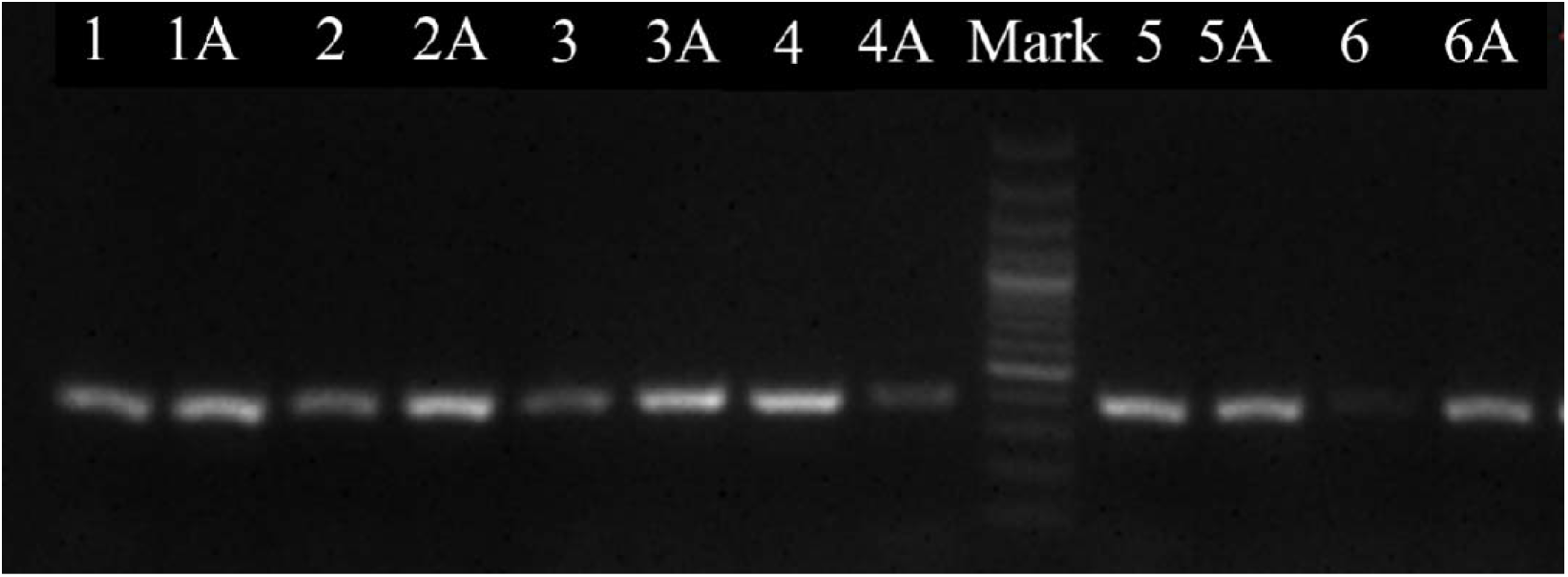
Results of gel electrophoresis of fecal sample. Samples with numeric labels (eg 1) correspond to DNA extracted from fresh fecal samples, while those with an “A” suffix (eg 1A) represent DNA extracted from the same individual fecal sample after six months of preservation. Specifically, 1 and 1A correspond to *Elaphodus cephalophus*, 2 and 2A to *Ailurus styani*, 3 and 3A to *Prionailurus bengalensis*, 4 and 4A to *Macaca thibetana*, 5 and 5A to *Hystrix brachyura* and 6 and 6A to *Sus scrofa*.

**FIGURE 4.**
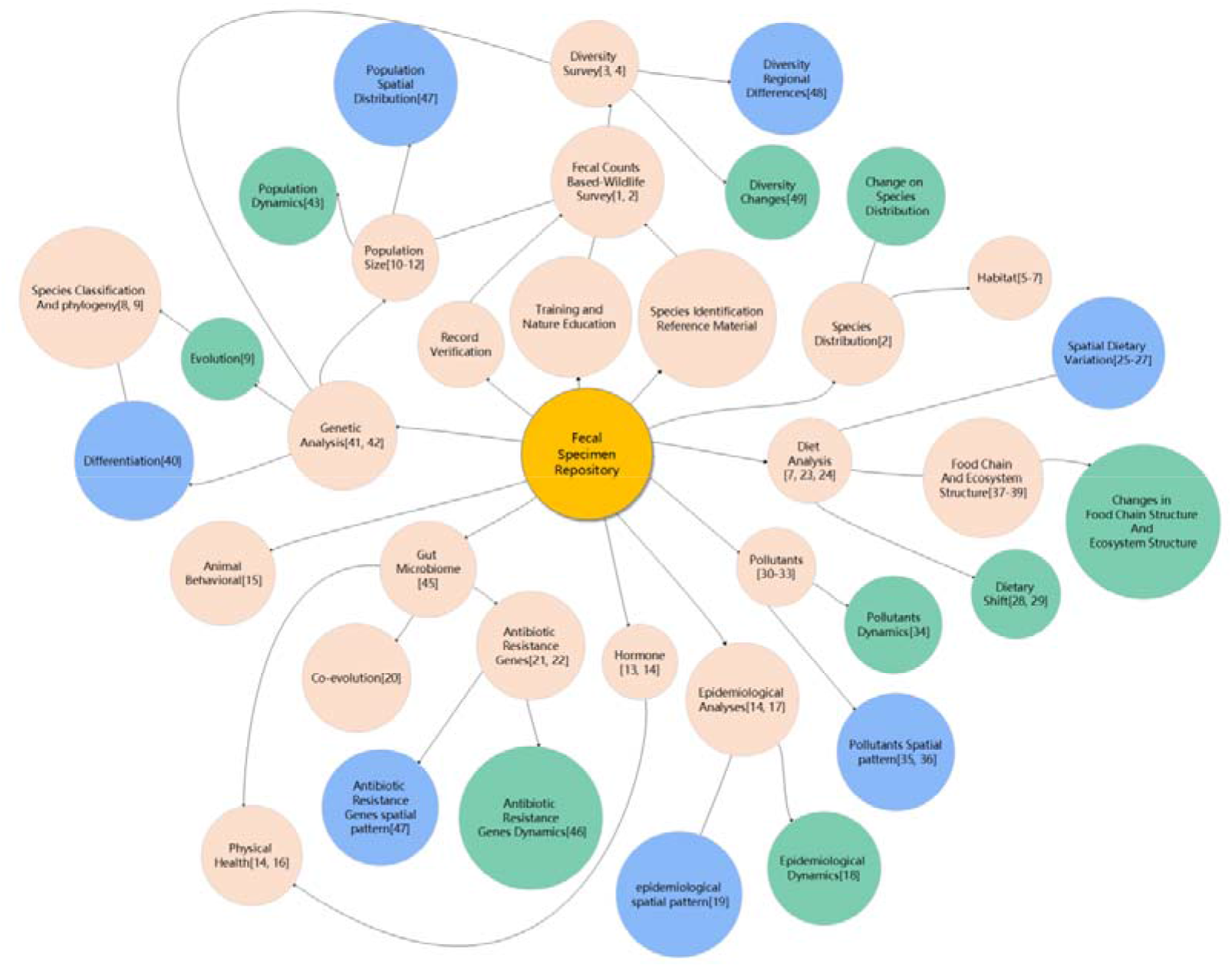
Existing and potential applications of a wildlife fecal specimen repository. Red circles indicate research and application directions, blue circles represent spatial expansion of studies, and green circles denote temporal expansion. The numbers in parentheses correspond to references in fecal research (Appendix 1).

Sequence homology was evaluated using BLAST against the NCBI database. Appendix Table 1 presents the sequence data and key metrics (total score, expected value, query cover, and percent identity) for fecal samples before and after six months. The consistently low expected values indicate high sequence reliability. All sequences exhibited a 97% identity, with *Prionailurus bengalensis* being the only exception, possibly due to sample degradation. These results confirm that DNA extracted from preserved specimens remains highly consistent with that from fresh samples, ensuring reliable species identification.

### Heavy Metal

Heavy metals were successfully detected in all fecal samples, demonstrating the applicability of fecal samples for heavy metal analysis (Table 3). Although concentrations varied among species, Cr, Cu, Zn, As, Cd, and Pb were consistently identified. In particular, the giant Panda (*Ailuropoda melanoleuca*) exhibited the highest Cr levels, while the Tufted Deer (*Elaphodus cephalophus*) had the highest As concentration.

**Table 3.**
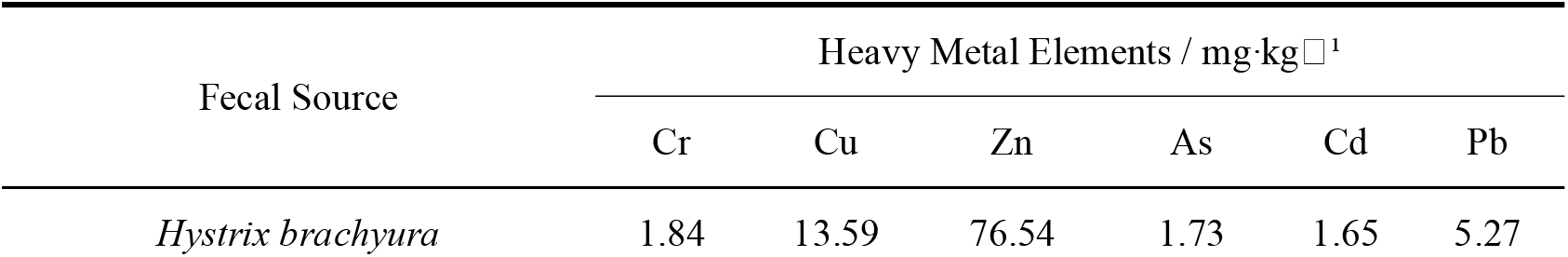
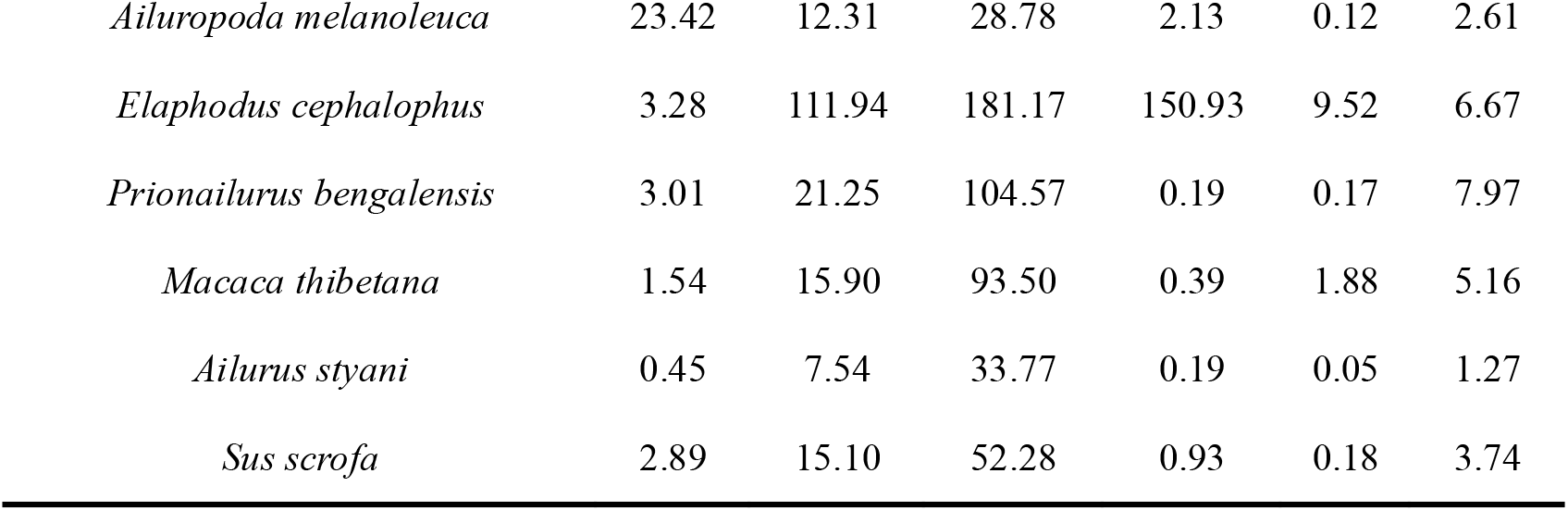
Heavy metal content in fecal specimens.

| Fecal Source | Heavy Metal Elements / mg·kg<sup>-1</sup> |  |  |  |  |  |
| --- | --- | --- | --- | --- | --- | --- |
|  | Cr | Cu | Zn | As | Cd | Pb |
| *Hystrix brachyura* | 1.84 | 13.59 | 76.54 | 1.73 | 1.65 | 5.27 |
| *Ailuropoda melanoleuca* | 23.42 | 12.31 | 28.78 | 2.13 | 0.12 | 2.61 |
| *Elaphodus cephalophus* | 3.28 | 111.94 | 181.17 | 150.93 | 9.52 | 6.67 |
| *Prionailurus bengalensis* | 3.01 | 21.25 | 104.57 | 0.19 | 0.17 | 7.97 |
| *Macaca thibetana* | 1.54 | 15.90 | 93.50 | 0.39 | 1.88 | 5.16 |
| *Ailurus styani* | 0.45 | 7.54 | 33.77 | 0.19 | 0.05 | 1.27 |
| *Sus scrofa* | 2.89 | 15.10 | 52.28 | 0.93 | 0.18 | 3.74 |

## Discussion

Wildlife feces are a widely used and highly informative noninvasive resource in ecological research, offering critical insights into various aspects such as species identification (Kurose et al. 2005; Walker et al. 2016), dietary composition (Green 1987; Węgrzyn et al. 2018), environmental contamination (Knapp et al. 2018; Andersson Stavridis et al. 2024), pathogen surveillance (Sinclair et al. 2008), and genetic diversity (Hu et al. 2018) (Figure 1). However, most fecal studies rely on fresh or short-term preserved samples, limiting their applicability for long-term ecological and conservation research. Here, we introduce a systematic and practical method for preparing fecal specimens that preserves both morphological integrity and biological and chemical composition over extended periods.

The fecal specimen preparation method used in this study effectively preserved the morphological characteristics of the samples. Compared to existing preservation methods, our approach maintains specimen stability in terms of size, color, and natural sheen while preserving delicate components such as hair and plant residues. Furthermore, hardness tests indicated strong mechanical properties, suggesting improved resistance to compression and mechanical damage, which is crucial for long-term storage and handling. In contrast, conventional preservation techniques have notable limitations. Alcohol fixation preserves genetic material, but compromises morphological integrity, hindering direct observation and measurement after immersion. Freezing slows degradation but is logistically challenging, costly for long-term storage, and prone to morphological damage due to repeated freeze-thaw cycles. Desiccation with silica gel, while simple, leaves specimens fragile during handling and susceptible to decomposition, mold growth, and insect damage over time.

Our experiments confirm that fecal specimens effectively preserve both the DNA and heavy metal composition. Even after six months of preservation, DNA was successfully extracted and the species identification results remained consistent with those obtained from fresh fecal samples. Heavy metals were also successfully detected in the preserved specimens, highlighting their potential for retrospective studies of environmental impacts on wildlife. However, since our study only evaluated DNA integrity over six months, more research is needed to assess its longer-term stability. For heavy metal analysis, an additional concern is the potential introduction of contaminants during specimen preparation and storage, which must be carefully controlled to ensure reliable concentration measurements and accurate retrospective assessments.

This method improves reliability by enabling repeated validation and reanalysis, thus ensuring research reproducibility, a fundamental requirement for advancing field-based animal ecology studies. Additionally, it facilitates long-term data archiving and supports retrospective studies, extending fecal sample applications across multiple disciplines, particularly in monitoring environmental and biodiversity changes under global change pressures.

## Perspective: Potential Applications

Wildlife feces contain rich biological information, making them valuable for ecological and conservation research. The preservation of fecal specimens in natural history museums and research collections allows long-term studies, facilitating comparative analyses across time and geographic regions, thus expanding their scientific applications across multiple research disciplines (Figure 1).

Fecal samples provide essential data for species distribution mapping (Jenkins & Manly 2008; Napolitano et al. 2008) and habitat dynamics analysis (Bashir et al. 2020; Sand et al. 2021; Seki et al. 2023). Long-term preservation ensures the integrity of diagnostic morphological traits, allowing repeated species verification and reducing misidentification. Curated fecal specimens serve as permanent reference materials for taxonomic comparison and species validation, which benefits both research and professional training. These specimens can aid in the training of field ecologists, conservation officers, and wildlife patrol teams, improving the accuracy of in-field species recognition.

Furthermore, preserved fecal samples support further molecular analyzes, such as DNA genotyping for species and individual identification, improving the reproducibility of the data in field-based ecological studies. By maintaining both morphological and genetic integrity over time, fecal specimen collections improve the reliability of long-term biodiversity assessments.

Fecal analysis is a widely used method for studying animal diets (Węgrzyn et al. 2018), providing insights into dietary composition, seasonal variation, and spatial differences (Suter et al. 2004; Stenset et al. 2016). Long-term preservation of specimens allows for cross-regional and historical dietary comparisons, offering a valuable perspective on shifting ecological conditions. For example, analyzing undigested food remains in snow leopard feces or identifying prey species through hair or DNA analysis allows researchers to assess prey composition and relative abundance, providing key insights into food resource availability (Shrestha et al. 2022). Establishing a repository of fecal specimens from wildlife would allow large-scale comparative diet studies in time and geography, improving our understanding of the feeding ecology of species, the use of resources, and the dynamics of the ecosystem.

Fecal specimens serve as sentinels for both infectious diseases and environmental contaminants (Levin et al. 2021). By detecting pathogens and toxins in feces, researchers can track the spatial and temporal dynamics of disease outbreaks (Gnat et al. 2015; Schilling et al. 2022). Unlike traditional disease monitoring, which relies on short-term field sampling, long-term fecal specimen archives allow retrospective epidemiological analyses and early detection of emerging pathogens.

Fecal samples also act as bioindicators of environmental contamination, reflecting the accumulation and fluctuation of pollutants within ecosystems (Knapp et al. 2018; Andersson Stavridis et al. 2024). A fecal specimen repository would provide an invaluable resource for long-term, large-scale environmental pollution monitoring, facilitating assessments of historical pollutant exposure and its effects on wildlife populations.

Beyond species identification, diet analysis, and environmental monitoring, a fecal specimen repository would support a wide range of interdisciplinary research applications. In microbiome research, preserved fecal samples offer opportunities to study long-term changes in gut microbial composition, contributing to our understanding of host physiology, host-microbe coevolution, and microbial antibiotic resistance (Pell 1997; Cenit et al. 2015; Menke et al. 2015; Guo et al. 2020; Wu et al. 2021). Genetic analysis of fecal DNA has advanced studies on evolutionary history, population size, genetic diversity, species classification, and phylogeny (Reed et al. 1997; Kohn et al. 1999; Joshi et al. 2020; Delibes-Mateos et al. 2023). In conservation science, fecal samples contribute to noninvasive population monitoring, allowing assessments of relative abundance and activity patterns (Ferretti et al. 2016; Davis et al. 2022). They also serve as critical tools for behavioral studies, offering information on species interactions, ecological adaptations, and movement patterns in natural habitats (Hegab et al. 2015).

By establishing a standardized fecal specimen repository, future research can leverage these collections for large-scale spatial and temporal comparative studies, allowing novel discoveries in ecology, conservation, and environmental sciences.

## Conclusions

Establishing a dedicated fecal specimen repository would bridge a critical gap in ecological and conservation research, transforming feces from a short-term diagnostic tool into a long-term resource for monitoring biodiversity. Such collections have the potential to become indispensable for species conservation, environmental assessment, and scientific education.

Although this study outlines key research applications, future advancements in molecular analysis, machine learning, and environmental forensics may unlock even broader and more innovative uses, further expanding the role of fecal specimens in ecology and global biodiversity research.

## Supporting information

Appendix 1 is provided to explain the figures in the main text.

Appendix Table 1 serves as supplementary data for the main text.

The title page contains details such as author information, acknowledgments, declarations of competing interests, and funding sources.

