## Appendix Table 1 serves as supplementary data for the main text. for "A Method for Preparing Morphologically Preserved Wildlife Fecal Specimens for Long-Term Ecological Studies"

| Fecal specimens | Timepoint | Sequence | Sequence length, bp | Total score | Query cover | Expect value | Percent identity | Accession number |
| --- | --- | --- | --- | --- | --- | --- | --- | --- |
| *Elaphodus cephalophus* | Before | CCATGGAGCTTTAATTAACTAATTCAAAAAGAAACTACTAACGACCCAACAGGAATAATATATCTCTTTTATGAATTAGCAATTTAGGTTGGGGCGACCTCGGAGGACAAAATAGCCTCCGAGTGATTATAAATCTAGACTTACCAGTCAAAATGCTTAATCACTTATTGATCCAAAAATTCTTTTGATCAACGGAACAAGTTACCCTAGGGATAACAGCGCAATCCTATCCGAGAGTCCATATCGACAATAGGGTTTACGACCTCGATGTTGGATCAGGACATCCTAATGGTGCAGCAGCTATTAAAGGTTCGTTTGTTCAACGATTAAAGTCCTACGTGATCTGAGTTCAGACCGGAGCAATCCAGGGGGG | 373 | 678 | 98% | 0 | 100.00% | OL702785.1 |
|  | After | TTGAGAAGAACCCCTTGGAGCTTTAACTACTTAGTCCAAAGAAATAAATTTTACTACCAAGGAAACAACAAGACTCTTTATGGACTAACAGCTTTGGTTGGGGTGACCTCGGAGAATAAAAAATCCTCCGAGCGATTTTAAAGACTAGACCTACAAGTCAAATCACACAATCGCTTATTGATCCAAAAAATTGATCAACGGAACAAGTTACCCTAGGGATAACAGCGCAATCCTATTCAAGAGTCCATATCGACAATAGGGTTTACGACCTCGATGTTGGATCAGGACATCCCGATGGTGCAACCGCTATCAAAGGTTCGTTTGTTCAACGATTAAAGTCCTACGTGATCTGAGTTCAGACCGGAGTAATCCAGTGGGGTTTTCTATTA | 389 | 673 | 99% | 0 | 97.95% | MN251783.1 |
| *Ailurus styani* | Before | TTGAGAAGATCCTATGGAGCTTCAATTAACTGACCCAATATAGATTAACCATATTTAACCAACCAGGGATATCATAACTCTATATCTGGGTTAGCAATTTAGGTTGGGGTGACCTCGGAGAAAAAACCATCCTCCGAGTGATACAATTCTAGACTTACTAGTCGAAACGCTCTATCATTTATTGATCCAAAATTTTTGATCAACGGAATAAGTTACCCTAGGGATAACAGCGCAATCCTATTCAAGAGTCCATATCAACAATAGGGTTTACGACCTCGATGTTGGATCAGGACATCCTAATGGTGCAGCAGCTATTAAGGGTTCGTTTGTTCAACGATTAAAGTCCTACGTGATCTGAGTTCAGACCGGAGCAATCCAGTGGGGTTATCTATTA | 394 | 688 | 99% | 0 | 98.23% | NC_009691.1 |
|  | After | TGAGAAGATCCCTTGGAGCTTCAATTAACTGACCCAATATAGATTAACCATATTTAACCAACCAGGGATATCATAACTCTATATCTGGGTTAGCAATTTAGGTTGGGGTGACCTCGGAGAAAAAACCATCCTCCGAGTGATACAATTCTAGACTTACTAGTCGAAACGCTCTATCATTTATTGATCCAAAATTTTTGATCAACGGAATAAGTTACCCTAGGGATAACAGCGCAATCCTATTCAAGAGTCCATATCAACAATAGGGTTTACGACCTCGATGTTGGATCAGGACATCCTAATGGTGCAGCAGCTATTAAGGGTTCGTTTGTTCAACGATTAAAGTCCTACGTGATCTGAGTTCAGACCGGAGCAATCCAGTGGGG | 383 | 680 | 98% | 0 | 99.21% | NC_009691.1 |
| *Prionailurus bengalensis* | Before | CGCCTGGATTACTCCGGTCTGAACTCAGATCACGTAGGACTTTAATCGTTGAACAAACGAACCTTTGATAGCTGCTGCACCATCGGGATGTCCTGATCCAACATCGAGGTCGTAAACCCTATTGTCGATATGGACTCTGAAATAGGATTGCGCTGTTATCCCTAGGGTAACTTGTTCCGTTGATCAAGTTWTTGGATCAATAARKGATGTAATAYTTTTGASTGKGTAGWMTWSATWTMAATCACTSGKAGGWKGTTKTGTTCTCCKAGGWCRCCCCAACMTAAATTGTCKGYTCAKSTAGAGGGTTGTTGTTCCTGTCGGGTATTTAGTGGAGTCTGTTTGAATCAATTAATTAAAGCTCCATAGGGGTCTTCTCA | 377 | 553 | 99% | 8.00E-153 | 91.18% | KX857783.1 |
|  | After | CTGGATTACTCCGGTCTGAACTCAGATCACGTAGGACTTTAATCGTTGAACAAACGAACCTTTGATAGCTGCTGCACCATCGGGATGTCCTGATCCAACATCGAGGTCGTAAACCCTATTGTCGATATGGACTCTGAAATAGGATTGCGCTGTTATCCCTAGGGTAACTTGTTCCGTTGATCAAGTTWTTGGATCAATARRKGATRTAATAYTTTTGWSTGKGTAGWMTWSATWTMAWTCACTSGKAGGWKGTTKTGTTCTSCKAGGWCRCCCCMACMTAAATTGTCKKTTCAKGTAGAGGKKTRTTGTWCSWGYCGGGTAGTTGTTGGAGACTGTTTGAATCAATTAATTAAAGCTCCATAGGGTCTTCTC | 372 | 501 | 100% | 3.00E-137 | 87.63% | KX857783.1 |
| *Macaca thibetana* | Before | TGAGAAGACCCCTATGGAGCTTTAATCTATTAATGCAATCAAAAACCAGATAAACCCACGGACCTCTAAACTACCGACACCTGCATTAAAAATTTTGGTTGGGGCGACCTCGGAGCACAACAAAACCTCCGAATAACAAATGCTAAGACCACACAAGTCAAAGCGAGCTAACACTCATAATTGATCCAATAATTTGATCAACGGAACAAGTTACCCTAGGGATAACAGCGCAATTCTATTCTAGAGTCCATATCGACAATAGAGCTTACGACCTCGATGTTGGATCAGGACATCCTAATGGTGCAGCAGCTATCAAGGGTTCGTTTGTTCAACGATTAAAGTCCTACGTGATCTGAGTTCAGACCGGAGCAATCCAGGCGGGGTTTCTATTA | 392 | 695 | 99% | 0 | 98.97% | NC_011519.1 |
|  | After | ACTGGATTGCTCCGGTCTGAACTCAGATCACGTAGGACTTTAATCGTTGAACAAACGAACCCTTGATAGCTGCTGCACCATTAGGATGTCCTGATCCAACATCGAGGTCGTAAGCTCTATTGTCGATATGGACTCTAGAATAGAATTGCGCTGTTATCCCTAGGGTAACTTGTTCCGTTGATCAAATTATTGGATCAATTATGAGTGTTAGCTCGCTTTGACTTGTGTGGTCTTAGCATTTGTTATTCGGAGGTTTTGTTGTGCTCCGAGGTCGCCCCAACCAAAATTTTTAATGCAGGTGTCGGTAGTTTAGAGGTCCGTGGGTTTATCTGGTTTTTGATTGCATTAATAGATTAAAGCTCC | 362 | 669 | 100% | 0 | 100.00% | NC_011519.1 |
| *Hystrix brachyura* | Before | CTTGAGAAGACCCTATGGAGCTTTAATTAATTTACATATACTAAAACACTTAACTTTCCTAAGGGTTACAAAACACTAGTACTATGTAAATAATTTTGGTTGGGGTGACCTCGGAGAACAAAAAAACCTCCGAATGATTTTAACCTAGACACTACAAGTCAAAGTCAAAAATCATCAATTGACCCAGTACAAACTGATCAACGAACCAAGTTACCCTAGGGATAACAGCGCAATCCTATTCTAGAGTTCTTATCGACAATAGGGTTTACGACCTCGATGTTGGATCAGGACTTCCCAATGGTGCAGCCGCTATTAAAGGTTCGTTTGT*CAACGATTAAAGTCCTACGTGATCTGAGTTCAGACCGGAGTAATCCAGGCGGGTTTCTATGAGGTCGGTTGAGATACCCCGCATCCAGGTCGGTTTCTATGA | 430 | 695 | 90% | 0 | 99.22% | NC_050263.1 |
|  | After | TGAGAAGACCCCCATGGAGCTTTAATTAATTTACATATACTAAAACACTTAACTTTCCTAAGGGTTACAAAACACTAGTACTATGTAAATAATTTTGGTTGGGGTGACCTCGGAGAACAAAAAAACCTCCGAATGATTTTAACCTAGACACTACAAGTCAAAGTCAAAAATCATCAATTGACCCAGTACAAACTGATCAACGAACCAAGTTACCCTAGGGATAACAGCGCAATCCTATTCTAGAGTTCTTATCGACAATAGGGTTTACGACCTCGATGTTGGATCAGGACTTCCCAATGGTGCAGCCGCTATTAAAGGTTCGTTTGTTCAACGATTAAAGTCCTACGTGATCTGAGTTCAGACCGGAGTAATCCAGGCGGGTTTTCTATT | 390 | 688 | 100% | 0 | 98.47% | NC_050263.1 |
| *Sus scrofa* | Before | GGGGGTATAGATCCGTAGGACTTTATCGTTGAACAAACGAACCTTTAATAGCGGTTGCACCATTTGGGTGTCCTGATCCAACATCGAGGTCGTAAACCCTATTGTCGATAGGAACTCTAGAATAGGATTGCGCTGTTATCCCTAGGGTAACTTGTTCCGTTGATCAAAATTTTGGATCAATAAGTGATGTTATGGTTATTTTGACTGGTTTGTCTAGATTAAAATCACTCGGAGGGTTTTTTGTACTCCGAGGTCACCCCAACCGAAATTGCTAGTCCATGTTAAGTTATGTTTTATCCCTTTGTGGTTGAATTGTTTAACTTTTGGAATAGTTAATTAAAGCTCCGTAGGGTCTTCTCA | 360 | 634 | 98% | 3.00E-177 | 99.15% | KC493607.1 |
|  | After | TTGAGAAGACCCTATGGAGCTTTAATTAACTATTCCAAAAGTTAAACAATTCAACCACAAAGGGATAAAACATAACTTAACATGGACTAGCAATTTCGGTTGGGGTGACCTCGGAGTACAAAAAACCCTCCGAGTGATTTTAATCTAGACAAACCAGTCAAAATAACCATAACATCACTTATTGATCCAAAATTTTGATCAACGGAACAAGTTACCCTAGGGATAACAGCGCAATCCTATTCTAGAGTTCCTATCGACAATAGGGTTTACGACCTCGATGTTGGATCAGGACACCCAAATGGTGCAACCGCTATTAAAGGTTCGTTTGTTCAACGATTAAAGTCCTACGTGATCTGAGTTCAGACCGGAGCAATCCAGGTGGGTTTTATATT | 392 | 704 | 99% | 0 | 99.48% | KC493607.1 |
| Note:"Before" refers to DNA sequences and BLAST results from fresh fecal samples, while "After" represents those obtained six months after the samples were preserved as fecal specimens. | | | | | | | | |
