## Supplementary material for "A Method for Preparing Morphologically Preserved Wildlife Fecal Specimens for Long-Term Ecological Studies": The title page contains details such as author information, acknowledgments, declarations of competing interests, and funding sources.

**Running Headline:**

Wildlife fecal specimen preparation method

**Corresponding Author:**
Qiang Dai
Chengdu Institute of Biology, Chinese Academy of Sciences
No. 23, Qunxian South Street, Chengdu 610041, China


**Acknowledgments:** We thank Mingxia Fu, Xinqiang Song, Xi Yang, Jun Kang, Sanchun Zhang, Xiaohong Zhang, and the staff of the Giant Panda Field Monitoring Team at the Yingjing Conservation and Management Station of Giant Panda National Park for their assistance during field sampling. This work was supported by the National Natural Science Foundation of China (Grant No. 32470536).

**Data Availability Statement**

The data that support the findings of this study are available from the corresponding author upon reasonable request.

**Conflict of Interest**

The authors declare no conflicts of interest related to this work.

**Author Contributions**

Jiahao Zhang and Qiang Dai conceived and designed the study. Jiahao Zhang, Xinrui Xu, and Yunqiao Zhang performed laboratory work. Dongling Zhang contributed to field sample collection and logistics. Qiang Dai supervised the project and revised the manuscript. All authors read and approved the final version of the manuscript.
